## Supplementary material for "Quantifying the effects of antibiotic resistance and within-host competition on strain fitness in *Streptococcus pneumoniae*": S1 Supplementary Information

### Note A: The Proportional Hazards assumption

We use survival analysis to analyse longitudinal data on pneumococcal carriage since (a) it can directly model time to event data (time to clearance or time to establishment), and (b) it can easily accommodate censorship in the data. Specifically, we use a survival regression method called the Cox Proportional Hazards model, which is the most generally applicable regression model. The idea of the Cox model is that the log-hazard (risk per unit time) of an event varies linearly with the covariates and the baseline hazard. The Cox model is non-parametric in that the baseline hazard function is not fixed unlike in other survival models.

The model relies on the proportional hazards assumption: that the ratio of hazards of a covariate between any two people is a time-independent constant. We test this for our basic Cox model where clearance rate is modelled as dependent on the serotype of the focal strain (Fig Ba). For other models, the time-varying covariates can capture the violations to the rule by accounting for changing covariate values. The test involves confirming that the inferred hazard coefficient (for instance, measured as the scaled Schoenfeld residuals of covariates) do not vary with time (Table A). We find that serotypes 14, 19A, 19B and 23F do not satisfy the proportional hazards test (at least one of the two p-values < 5%). However, this would not be a limitation for our results since some proportional hazard violations are expected given the large size of our dataset and number of covariates included.

**Table A.** Testing the proportional Hazards assumption: Two test statistics are computed for each serotype - (1) KM (Kaplan meier) that checks for violations of the proportional hazards assumption by comparing the survival curves of different groups (stratified by a covariate) to see if they are parallel over time, and (2) Rank based test that evaluates the proportional hazards assumption by looking at the correlation between time and the scaled Schoenfeld residuals of the model

| Variable | Test | Test Statistic | p-value | -log2(p) |
|---|---|---|---|---|
| 1 | km | 0.17 | 0.68 | 0.56 |
| | rank | 3.40 | 0.07 | 3.94 |
| 10A | km | 0.44 | 0.51 | 0.98 |
| | rank | 2.60 | 0.11 | 3.23 |
| 10B | km | 0.02 | 0.89 | 0.17 |
| | rank | 1.34 | 0.25 | 2.01 |
| 10F | km | 0.49 | 0.48 | 1.05 |
| | rank | 0.67 | 0.41 | 1.28 |
| 11A | km | 0.06 | 0.80 | 0.32 |
| | rank | 0.38 | 0.54 | 0.90 |
| 12F | km | 0.08 | 0.78 | 0.35 |
| | rank | 0.15 | 0.70 | 0.52 |
| 13 | km | 0.94 | 0.33 | 1.59 |
| | rank | 0.31 | 0.58 | 0.79 |
| 14 | km | 0.38 | 0.54 | 0.90 |
| | rank | 5.26 | 0.02 | 5.52 |
| 15A | km | 2.21 | 0.14 | 2.87 |
| | rank | 0.60 | 0.44 | 1.18 |
| 15B_C | km | 1.39 | 0.24 | 2.07 |
| | rank | 1.56 | 0.21 | 2.24 |
| 16F | km | 2.12 | 0.14 | 2.79 |
| | rank | 0.88 | 0.35 | 1.52 |
| 17F | km | 0.02 | 0.89 | 0.18 |
| | rank | 2.34 | 0.13 | 2.99 |
| 19A | km | 4.33 | 0.04 | 4.74 |
| | rank | 0.01 | 0.90 | 0.15 |
| 19B | km | 6.35 | 0.01 | 6.41 |
| | rank | 0.83 | 0.36 | 1.47 |
| 19F | km | 2.64 | 0.10 | 3.27 |

| Variable | Test | Test Statistic | p-value | -log2(p) |
|---|---|---|---|---|
| | rank | 2.35 | 0.13 | 3.00 |
| 20 | km | 0.23 | 0.64 | 0.66 |
| | rank | 0.18 | 0.67 | 0.58 |
| 21 | km | 0.30 | 0.58 | 0.78 |
| | rank | 0.97 | 0.32 | 1.63 |
| 22A | km | 0.00 | 0.94 | 0.08 |
| | rank | 1.66 | 0.20 | 2.34 |
| 22F | km | 0.53 | 0.47 | 1.10 |
| | rank | 0.04 | 0.85 | 0.24 |
| 23A | km | 0.12 | 0.73 | 0.45 |
| | rank | 0.15 | 0.70 | 0.51 |
| 23B | km | 0.15 | 0.70 | 0.52 |
| | rank | 0.50 | 0.48 | 1.05 |
| 23F | km | 3.37 | 0.07 | 3.91 |
| | rank | 5.72 | 0.02 | 5.90 |
| 24F | km | 0.06 | 0.81 | 0.31 |
| | rank | 1.27 | 0.26 | 1.94 |
| 28F | km | 0.61 | 0.43 | 1.21 |
| | rank | 0.99 | 0.32 | 1.65 |
| 29 | km | 0.08 | 0.78 | 0.35 |
| | rank | 0.02 | 0.88 | 0.19 |
| 3 | km | 0.59 | 0.44 | 1.17 |
| | rank | 0.25 | 0.62 | 0.70 |
| 33B | km | 0.04 | 0.84 | 0.25 |
| | rank | 2.56 | 0.11 | 3.19 |
| 33C | km | 0.09 | 0.76 | 0.39 |
| | rank | 2.68 | 0.10 | 3.30 |
| 33F | km | 0.30 | 0.89 | 0.17 |
| | rank | 0.25 | 0.62 | 0.70 |
| 34 | km | 0.30 | 0.58 | 0.78 |
| | rank | 0.38 | 0.54 | 0.90 |
| 35C | km | 1.23 | 0.27 | 1.90 |
| | rank | 0.93 | 0.33 | 1.58 |
| 35F | km | 2.26 | 0.13 | 2.91 |
| | rank | 0.88 | 0.35 | 1.53 |
| 38 | km | 0.28 | 0.60 | 0.74 |
| | rank | 0.00 | 0.98 | 0.02 |
| 4 | km | 0.21 | 0.65 | 0.62 |
| | rank | 1.06 | 0.30 | 1.72 |
| 45 | km | 0.00 | 0.98 | 0.03 |
| | rank | 1.03 | 0.31 | 1.69 |
| 46 | km | 0.01 | 0.94 | 0.09 |
| | rank | 1.06 | 0.30 | 1.72 |
| 5 | km | 0.02 | 0.90 | 0.15 |
| | rank | 0.09 | 0.76 | 0.39 |
| 6A_C | km | 3.36 | 0.07 | 3.90 |
| | rank | 0.03 | 0.87 | 0.20 |
| 6B | km | 1.45 | 0.23 | 2.13 |
| | rank | 0.50 | 0.48 | 1.06 |
| 7B | km | 1.71 | 0.19 | 2.39 |

| Variable | Test | Test Statistic | p-value | -log2(p) |
|---|---|---|---|---|
| | rank | 0.02 | 0.90 | 0.15 |
| 7F | km | 0.05 | 0.83 | 0.27 |
| | rank | 1.05 | 0.30 | 1.72 |
| 8 | km | 0.06 | 0.81 | 0.30 |
| | rank | 0.12 | 0.73 | 0.46 |
| 9L | km | 0.00 | 0.97 | 0.04 |
| | rank | 0.00 | 0.96 | 0.06 |
| 9N | km | 3.50 | 0.06 | 4.02 |
| | rank | 0.58 | 0.44 | 1.17 |
| 9V | km | 0.53 | 0.46 | 1.11 |
| | rank | 0.21 | 0.65 | 0.63 |

### Note B: Sensitivity to assumptions about start and end of carriage

Since exact dates of colonisation and clearance of strains cannot be known, we assumed that the colonisation or clearance event happened mid-way between the two sampling points where the potential event occurred. Here, we test the sensitivity of our results on within-host competition to this assumption (Fig Ca). If the sampling data indicates the start of carriage between $T_0$ and $T_1$, we take the time of establishment to be at $T_1$ - k*($T_1$ - $T_0$), where k=0.25 (Fig Ca - Case A) or 0.75 (Fig Ca - Case B). k=0.5 corresponds to results from the main text. Similarly if clearance occurs between times $T_2$ and $T_3$, then the time of establishment is $T_2$ + k*($T_3$ - $T_2$). We analyse the results on within-host competition for these values of k.

We find that the presence of within-host competitors always increases the rate of clearance of strains for k values of 0.25, 0.5 and 0.75 (Fig Cb(i)). The hazard coefficients of presence-of-competitor are 0.28 (CI=0.22 to 0.36) for k=0.25, 0.28 (CI=0.21 to 0.35) for k=0.5 and 0.25 (CI=0.18 to 0.32) for k=0.75. The effect of serotype on within-host competition is tested using a model with interaction coefficients between serotype and presence-of-competitor, and regressing the serotype coefficients with the interaction coefficients (Fig 3B). The slopes of this regression are always negative for k=0.25 to 0.75 (Fig Cb(ii)). The slope is -0.41 (CI=-0.60 to -0.21) for k=0.25, -0.40 (CI=-0.60 to -0.20) for k=0.5 and -0.42 (CI=-0.60 to -0.23) for k=0.75. Similarly, the effect of presence of competitors on rate of establishment is also not sensitive to the values of k tested (Fig Cb(iv)). The hazard coefficient of presence-of-competitor on establishment are -0.80 (CI=-0.90 to -0.69) for k=0.25, -0.79 (CI=-0.90 to -0.68) for k=0.5 and -0.75 ((CI=-0.86 to -0.64) for k=0.75.

The priority effect that we observe on the clearance of co-colonised strains is sensitive to the values of k tested (Fig Cb(iii)). The results of this analysis are significant for k=0.5 and 0.75. The difference in clearance rates between a resident and non-resident competitor is not significantly different from each other when k=0.25, but both have a positive hazard of clearance. The absence of an effect for k=0.25 might be driven by scenarios where there is consecutive loss and gain of different serotypes in a host (i.e., serotype replacement). When two different serotypes are detected in two consecutive samples in the data, then k=0.25 (and not 0.5 and 0.75) corresponds to the assumption that the earlier serotype is lost before establishment of the new serotype such that no co-colonisation occurs. In the case of k=0.5 and 0.75, the new serotype would be assumed to replace the pre-existing serotype shortly after co-colonisation. So the effect of a resident serotype may be acting primarily on invading strains that result in serotype replacement.

### Note C: Sensitivity to assumptions about serotype detection during sampling

In the analysis, we assume all serotypes present in samples are always observed. However, the sampling methods used to collect the data may fail to detect the presence of a serotype, especially when serotypes are present in low frequency in a host. Here, we perform a sensitivity analysis to test the effects of undersampling of serotypes on our results.

Our dataset consists of over 19,000 nasopharyngeal samples. To simulate undersampling, we remove observations (i.e. a serotype detected in a sample) with probability p. We regenerate data by simulating this undersampling step across all samples, for a range of values of p from 1% to 50%. We simulate a uniform non-detection rate across all samples, irrespective of within-host frequencies.

The results of the sensitivity analyses are shown in Fig D. The estimates for p=0 correspond to the effects of within-host competition we discuss in Fig 3 in the main text. The estimates for non-zero values of undersampling probability are shown for 10 random iterations. We find that the estimated effect of within-host competitors on clearance and establishment rates become weaker with undersampling (both in magnitude and in uncertainty), but are not qualitatively sensitive to the undersampling probabilities up to 50% (Fig Da and Dd). The priority effect - the difference in the coefficients between *presence-of-invader-competitor* and *presence-of-resident-competitor* - increases with undersampling (Fig Dc). This suggests that the estimated effect may diminish with greater sampling depth, and more complete data is required to confirm whether the effect we observe reflects a true priority effect. The uncertainty in the estimation of the trade-off between clearance during single-carriage and co-carriage in Fig Db increases with undersampling. This may be because increased undersampling decreases the number of serotypes in the dataset, and thereby decreases our power to infer serotype-specific differences in clearance rates. Overall, this analysis suggests the effects of within-host competition on establishment and clearance are robust to undersampling, but reduces our confidence in the priority effect.

### Note D: Effect of antibiotics on establishment rates

In the main text, we analysed the effect of antibiotic consumption on clearance rates of resistant and susceptible strains. Here, we report the effects of antibiotic consumption on establishment rates. Since comparison of establishment rates of resistant versus susceptible strains is confounded by the total prevalence of resistance in the population, we can only infer the effects of antibiotics on susceptible strains, captured by the variable *effective-drug-use*.

Similar to the corresponding clearance model, we perform survival analysis on the times to establishment of serotypes using *presence-of-competitor*, *effective-drug-use*, and *ineffective-drug-use* as covariates, and age of infant and serotype as stratifying variables. The hazard coefficient of *effective-drug-use* is 0.00 (CI=-0.09 to 0.09) (Fig Fa). We find no signal for an effect of drug consumption on the rate of establishment of drug-susceptible strains, even when analysed for amoxicillin / ampicillin separately (Fig Fc). However, the coefficient value of *effective-drug-use* is sensitive to the assumption of length of drug-use efficacy of 7 days, and *effective-drug-use* is associated with a positive value when this length is assumed to be 15 days (more in Note G).

### Note E: Effect of different antibiotics on clearance rates

In this section, we quantify how antibiotic usage (captured by the effect sizes of *effective-drug-use* and *ineffective-drug-use*) affects clearance rates across all drug treatments in the dataset.

We use a Cox proportional hazards model with serotype, *presence-of-competitor*, *effective-drug-use*, and *ineffective-drug-use* included as covariates (similar to the model in main text). The covariates *effective-drug-use*, and *ineffective-drug-use* are defined with respect to a single antibiotic at a time. The results of this analysis are shown in Fig Fb. Only four antibiotics are shown here; the model does not converge well for the others since there are only few recorded instances of use of those drugs. For amoxicillin/ampicillin, the hazard coefficient of *effective-drug-use* is 0.55 (CI=0.41 to 0.68) and of *ineffective-drug-use* is 0.34 (CI=0.13 to 0.54). This corresponds to an increase in clearance rate by 73% (hazard ratio of 1.73, CI=1.51 to 1.97) of susceptible strains and by 40% (hazard ratio of 1.40, CI=1.14 to 1.73) of resistant strains. Similar to the analysis across all drugs, we see that susceptible strains are cleared faster by amoxicillin/ampicillin than resistant strains. For all other antibiotics, 95% confidence intervals of their effect sizes overlap zero.

### Note F: Antibiotic-blind analyses of the costs of resistance

In the main text, we quantified the fitness costs of resistance, i.e., the effect of resistance on the clearance rate of strains under no antibiotic exposure conditions (using the covariate *resistance-without-drug*). Here, we look at whether resistance has any overall fitness effects *without* stratifying the data by drug consumption. In other words, we quantify how *resistance* affects fitness of a strain while being blind to whether the strain experiences antibiotic pressure. We test this for three fitness components - clearance and establishment rates, and within-host competition between strains.

First, we test whether resistant strains have a higher clearance rate in our dataset than susceptible strains. We run a time-varying proportional hazards model that includes serotype of focal strain, *presence-of-competitor* and resistance status of focal strain as covariates. The resistance variable is used in two ways (in separate analyses): (1) Resistance to antibiotic X, for all X, and (2) Resistance to any antibiotic. We do not include ceftriaxone resistance, since there are only two episodes of colonisation in the data that are ceftriaxone resistant. We find that the hazard coefficient of resistance is variable from positive to negative depending on the antibiotic tested (Fig Ga). For 3 antibiotics, the hazard is negative and significant, and for 2 antibiotics, is negative with CI overlapping zero. For these five antibiotics, the results suggest that resistance is associated with a longer duration of carriage. We find a signal indicating a clearance cost in resistance to chloramphenicol, which has a significant positive hazard of 0.24 (CI=0.02 to 0.47). This corresponds to a 27.4% higher clearance rate of chloramphenicol-resistant strains compared to chloramphenicol-susceptible strains.

We then look at the interaction term between *presence-of-competitor* and resistance in the above model, for all seven antibiotics. The interaction coefficients measure the hazard of clearance due to resistance (compared to susceptibility) in the presence of within-host competitors, thereby capturing the cost of resistance on within-host competition. None of the interaction terms between resistance and *presence-of-competitor* are significantly different from zero (Fig Gb). In this analysis, we find no signal for a cost of resistance on within-host competitive ability during clearance.

In the above analyses, we looked at how resistance of the *focal strain* affects its clearance. Next, we analyse how resistance status of a within-host competitor affects clearance of the focal strain. To do this, we define the time-dependent variables *presence-of-resist-competitor* and *presence-of-sens-competitor*, which indicate whether the within-host competitor, if present, is resistant or susceptible respectively. We use a time-varying proportional hazards model with the following covariates: serotype of focal strain, *presence-of-resist-competitor*, and *presence-of-sens-competitor*. The difference in the values of coefficients between *presence-of-resist-competitor* and *presence-of-sens-competitor*, signifies the effect of resistance on a competitor's ability to induce clearance of the focal strain. As before, we repeat the analysis for all seven antibiotics independently, and by assuming

| Figure | Model | AIC | AIC - min(AIC) |
|---|---|---|---|
| Fig 3A | Model (in main text) | 61620.21 | 0 |
| | Model without serotype | 63042.06 | 1421.85 |
| Fig 3B | Model | 89897.86 | 0 |
| Fig 3C | Model | 61491.22 | 0 |
| | Model without serotype | 63006.52 | 1515.30 |
| Fig 3D | Model | 13315.5 | 0 |
| Fig 5A | Model | 62180.7 | 0 |
| | Model without serotype | 62976.80 | 796.10 |
| | Model without competitor | 62852.89 | 672.2 |
| | Model - both | 63040.70 | 860.0 |
| Fig 5B | Model | 62162.50 | 0 |
| | Model without serotype | 62937.9 | 775.40 |
| Fig 5C | Model | 13329.30 | 0 |

**Table B.** Partial AIC values for Cox models from analyses in the main text.

resistance to *any* antibiotic. The hazard coefficients of *presence-of-resist-competitor* and *presence-of-sens-competitor* are not significantly different from each other for any of the antibiotics tested (Fig Gc). We do not find any signal for an effect of within-host competitors' resistance on the clearance rate of the focal strain.

Finally, we analyse the effect of resistance on establishment rate of strains. We test whether the competitive effect faced by an establishing strain depends on whether the resident strain is resistant or susceptible to antibiotics. This is achieved using survival analysis on 'time to establishment' data using the following covariates: *presence-of-resist-competitor* and *presence-of-sens-competitor*, with age of host and serotype of incoming strain used to stratify the baseline hazard rate. We find that the hazard coefficients associated with *presence-of-resist-competitor* are significantly lower than those of *presence-of-sens-competitor* (Fig H) for all resistances tested. This indicates an association of antibiotic-resistant resident strains with increased prevention of new establishment compared to susceptible strains. Such an effect could be driven by an association between antibiotic usage and carriage of resistant strains, such that the lower establishment rate is caused by antibiotic exposure. Overall, we do not find evidence for a cost of resistance on establishment rates.

### Note G: Assumptions about drug consumption in hosts

We assumed in our analyses that drug consumption, where included, has an effect in hosts for 7 days. We test the sensitivity of our results to this assumption by varying the assumed duration of drug efficacy to 5, 10 and 15 days (Fig J).

Antibiotic consumption is always associated with an increased rate of clearance of resistant and sensitive strains for all drug-use lengths tested (positive coefficients of *effective-* and *ineffective-drug-use* in Figs 5A and Ja). The difference between the two hazard coefficients however is sensitive to this assumption: the hazard of *effective-drug-use* is significantly higher than that of *ineffective-drug-use* when the drug-use duration is 10 days or higher. At drug-use length of 10 days, the hazard coefficients are 0.62 (CI=0.52 to 0.72) for *effective-drug*-use and 0.19 (CI=0.02 to 0.38) for *ineffective-drug-use*. The difference between these two coefficients signify the fitness benefit of resistance on rate of clearance of strains. Resistance is associated with a significant fitness benefit during antibiotic use when we assume the length of drug-use to be 10 or more days.

The result on the effect of drug use on rate of establishment is sensitive to the lengths of drug-use tested (Figs Fa and Jb). The strength of the coefficient of *effective-drug-use* is positive and significant for the case of 15 days, which corresponds to a higher establishment rate of sensitive strains. This may be because, by assuming 15 days, the model captures more cases of post-antibiotic-use establishment of sensitive strains. If so, then this coefficient captures the post-treatment repopulation of the host niche rather than the effect of treatment itself.

All results corresponding to measuring the costs of resistance are non-sensitive to the assumptions about length of drug-use. These include - the effect of resistance on clearance rate (Fig Jc), cost of resistance on competitive ability during clearance (Fig Jc) and establishment (Fig Jd), and the costs of resistance measured for individual resistances (Fig Je and Jf).

### Note H: Potential within-host gain or loss of resistance

In our analysis, we assume that resistance to antibiotics is not gained or lost during an episode of carriage, for instance via horizontal gene acquisition. One way to check for potential within-host resistance switches using our data is to look at episodes of carriage where we have multiple resistance measurements. In around 1.9% of all episodes of carriage where resistance status is known, measured resistance changed corresponding to either an S→R (0.9% cases) or R→S (1% cases) change. Some of these may be within-host resistance gain or loss if they also correspond to co-carriage with another strain.

We show that these cases do not affect the main results of our study by removing the 1.9% cases where discordant resistance measurements are observed and repeating our analysis. These results are shown in Fig K. Most of our results on the benefit and cost of resistance are not qualitatively affected by this reduction of data except in one case - corresponding to (Fig Kd) - where the significance of the coefficient is reduced. The hazard ratio corresponding to the within-host competitive cost of erythromycin resistance changed from 1.17 (CI=1.01 to 1.36) to 1.15(CI=0.99 to 1.34).

### Note I: Model selection and parsimony estimations

We use Akaike Information criterion (AIC) to calculate model parsimony. AIC is a measure of relative quality of statistical models that balances the risk of over-fitting by adding too many variables versus the risk of under-fitting by using an oversimplified model.

For a model with k independent variables and a maximum likelihood estimate of L, the value of AIC is given by the equation:

$$AIC = 2(k - ln(L)) \tag{A}$$

For each model, we extracted the AIC of the model, and compared to that of all models with a subset of covariates that can be used to analyse the effects we study in the model. In Cox models, we calculate the partial AIC since the model maximises a partial likelihood function that focuses on the relative hazard rates between individuals without requiring knowledge of the baseline hazard function. A lower partial AIC corresponds to a more parsimonious model. The values of partial AIC tested for different models are listed in Table B.

### Figures

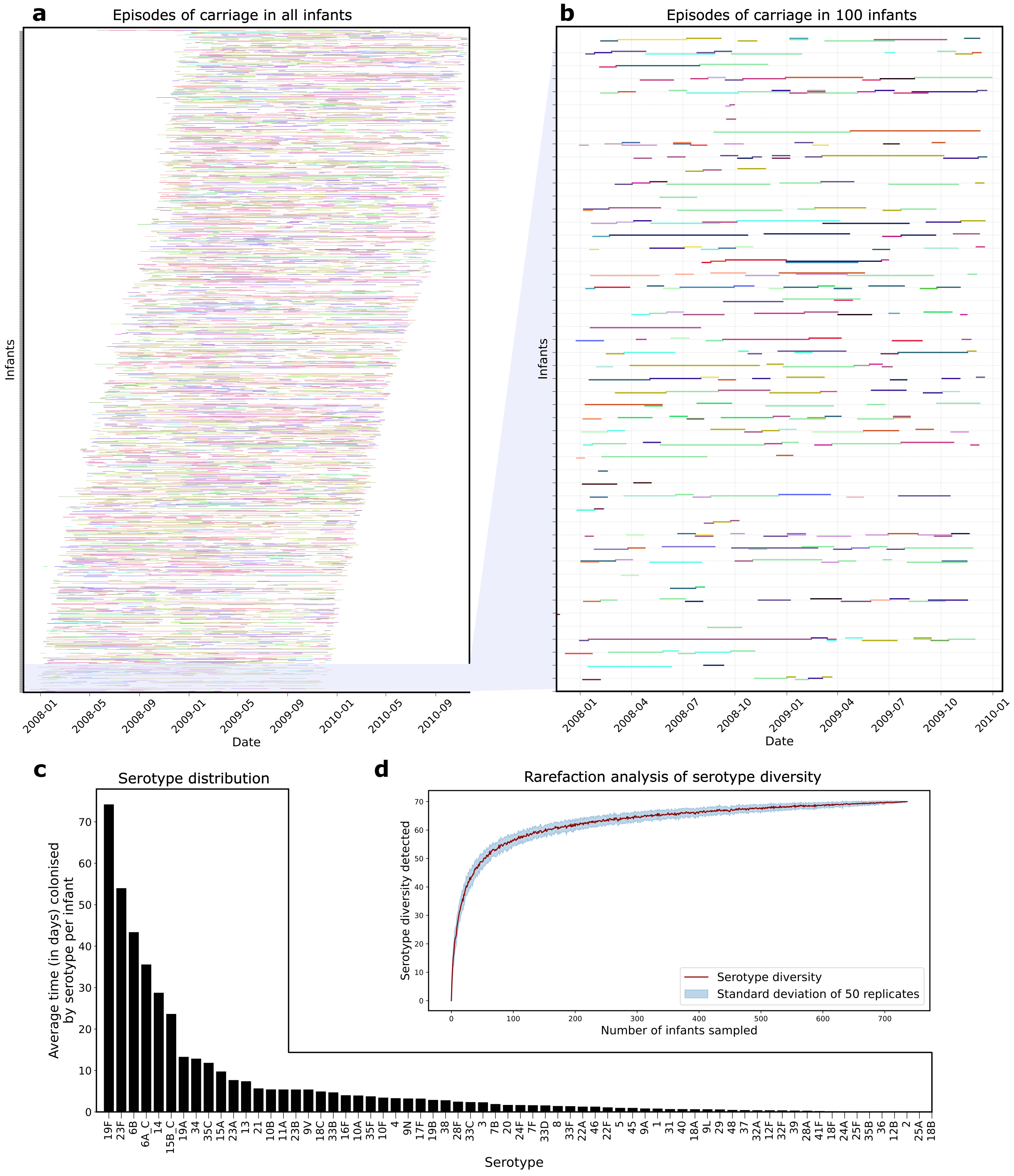


**Fig A.** Brief overview of the data used to run the survival analyses. (a) The durations of carriage of all observed strains in all infants used for the analyses. Each row along the vertical axis corresponds to an individual infant. Each coloured horizontal line represents the carriage of a strain in the host along time (shown on the X axis). The colours represent unique serotypes. The serotypes in each row are slightly offset vertically to aid the visualization of overlapping carriages within an infant. (b) The first 50 infants (from the bottom of subplot a) are zoomed into to show carriage of strains in individual infants. (c) Serotype distribution in the dataset. Each bar corresponds to the sum of all observed durations of carriage of the serotype divided by the total number of infants. (d) Rarefaction curve that describes increasing detection rate of serotypes with increasing sampling rate of infants in the dataset. Around 60 out of 68 serotypes are detected with a sample size of just 200 infants. The data underlying this figure can be found in https://doi.org/10.5281/zenodo.15800560

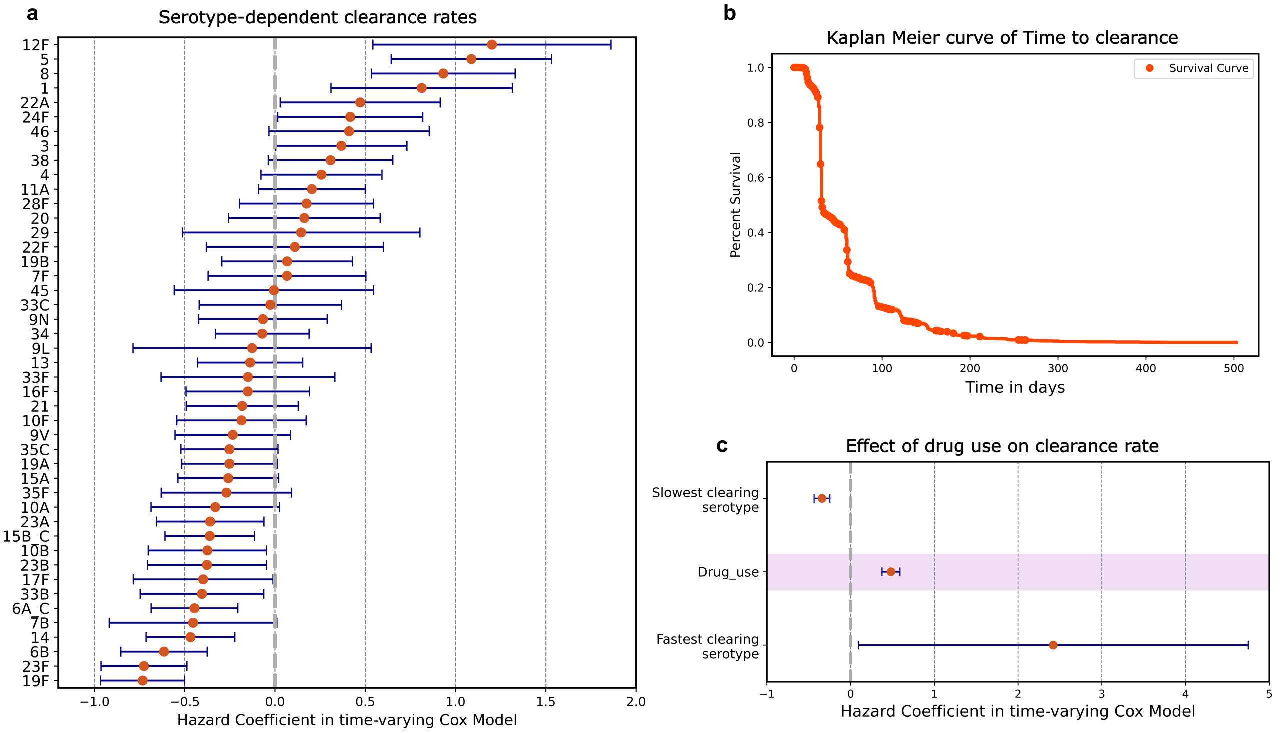


**Fig B.** Basic survival models to describe pneumococcal dynamics (a) Clearance rate (and thus the duration of carriage) varies with the serotype of the strain. (b) For the model in a, the survival curve that summarises the duration of carriage of all strains in our data. On the Y axis is plotted the fraction of episodes of carriage that have a duration of carriage that lasts at least as long as the value corresponding to the X axis. (c) Does antibiotic consumption affect the clearance rate of strains? The Drug use covariate has a positive coefficient of 0.46 (CI=0.36 to 0.57) indicating increased clearance rates associated with antibiotic use. The data underlying this figure can be found in https://doi.org/10.5281/zenodo.15800560

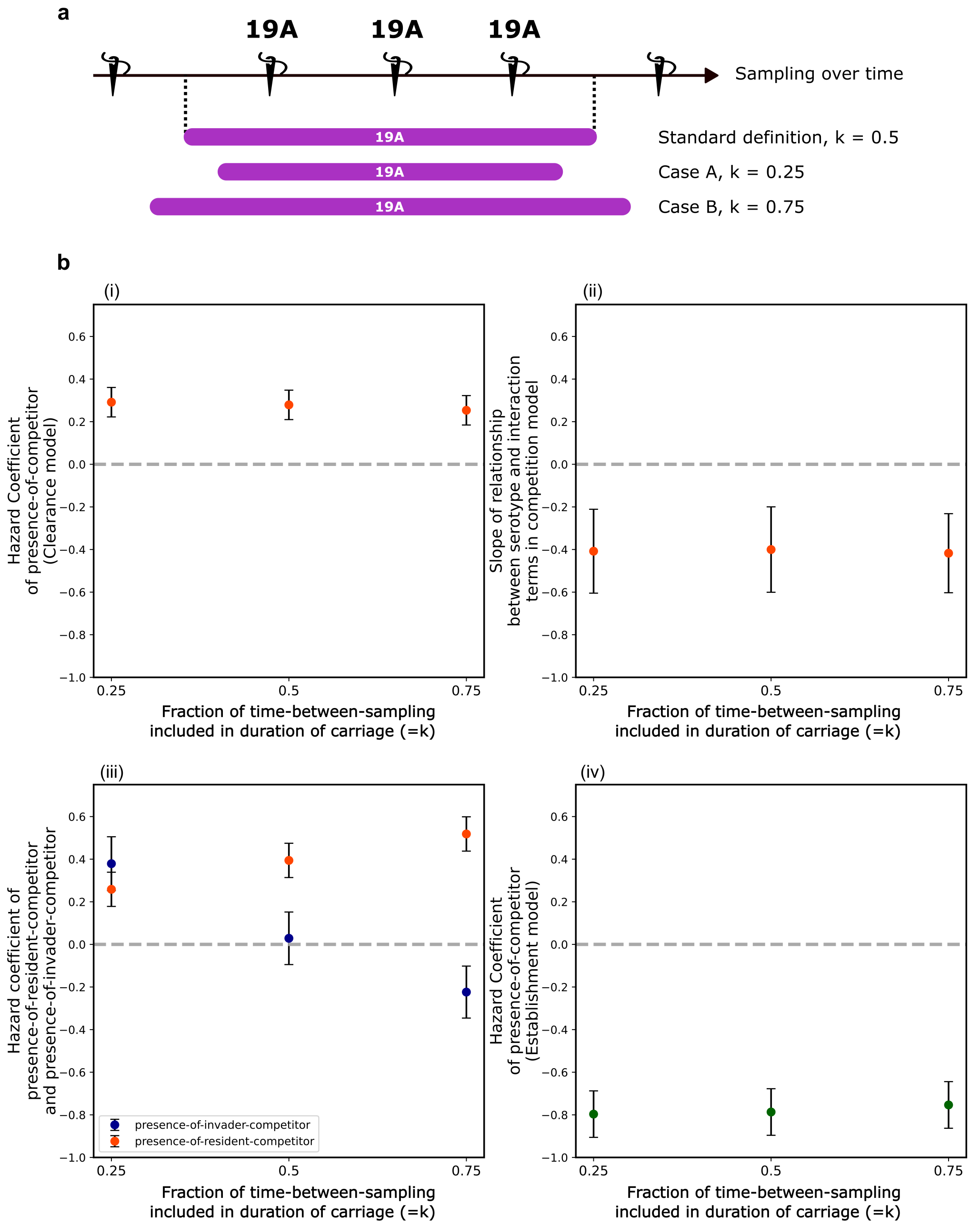


**Fig C.** Testing sensitivity to assumptions about start and end of carriage. (a) An example schematic is shown where a strain with serotype 19A is detected in 3 consecutive samples. The standard definition of duration of carriage assumes start and end of carriage to be half-way between two sampling dates. We modify this assumption in Case A and B where these start and end dates are taken to be 25% or 75% of the duration between two sampling dates respectively. (b) We test the effects of assumptions in (a) on the results on within-host competition. The effects of within-host competition on clearance and establishment are not sensitive to the assumption about start and end dates of carriage, except in one case. The resident-priority effect in clearance (shown in Figure 2C) between co-colonised strains is not significant in the case of 25% as seen in b(iii). The data underlying this figure can be found in https://doi.org/10.5281/zenodo.15800560

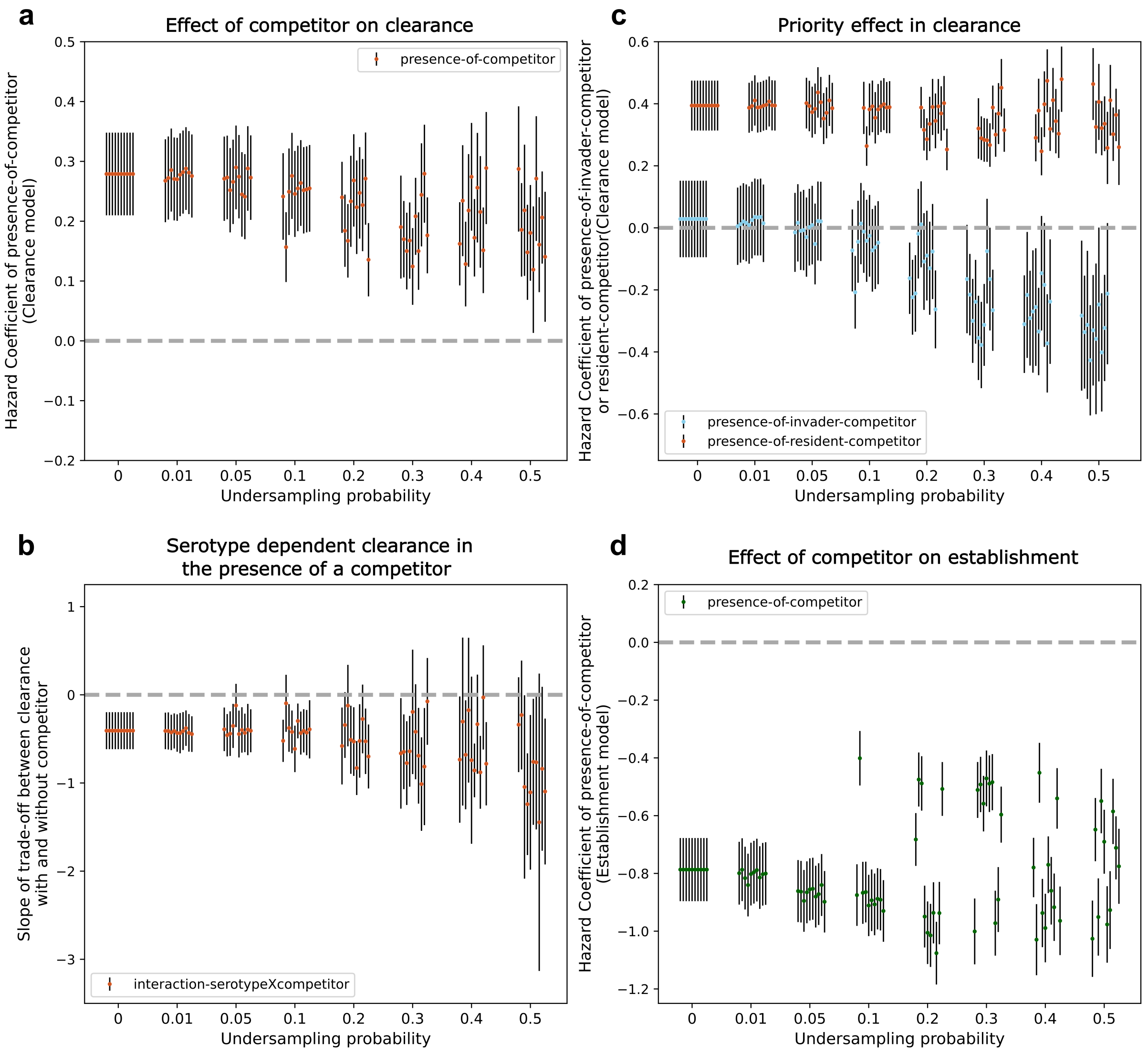


**Fig D.** Testing sensitivity of results on within-host competition to undersampling of data. a-d corresponds to the results from Figure 2 A-D. Each estimate is repeated 10 times - shown by the jittered error bars. All error bars reflect 95% confidence intervals. (a) The effect of competition on increasing the rate of clearance. (b) The negative correlation between hazard coefficients of serotypes and those of serotype-competitor interaction terms. (c) The difference in clearance risks between a resident competitor and an invader competitor. (d) The effect of competition on reducing the rate of establishment of new strains. In all plots, the estimates corresponding to probability of zero corresponds to the estimates from the main text. The data underlying this figure can be found in https://doi.org/10.5281/zenodo.15800560

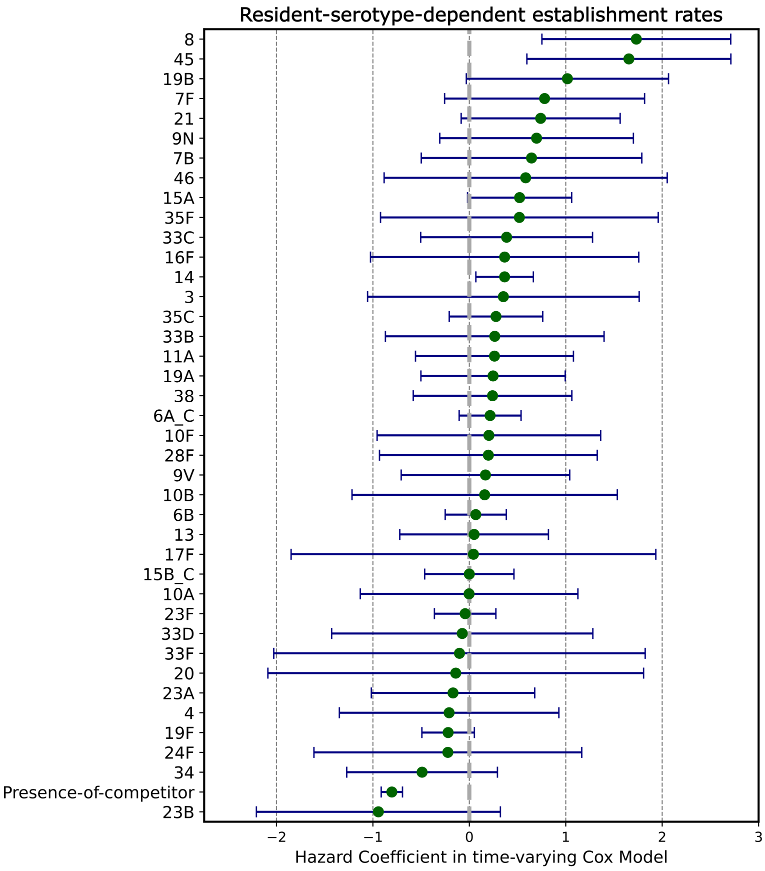


**Fig E.** Effect of serotype of the resident strain on the rate of establishment of incoming strains. The model includes resident serotypes as covariates to look at their effects on establishment. Serotypes can have a range of different competitive abilities, ranging from preventing new establishment to facilitating new establishment. The data underlying this figure can be found in https://doi.org/10.5281/zenodo.15800560

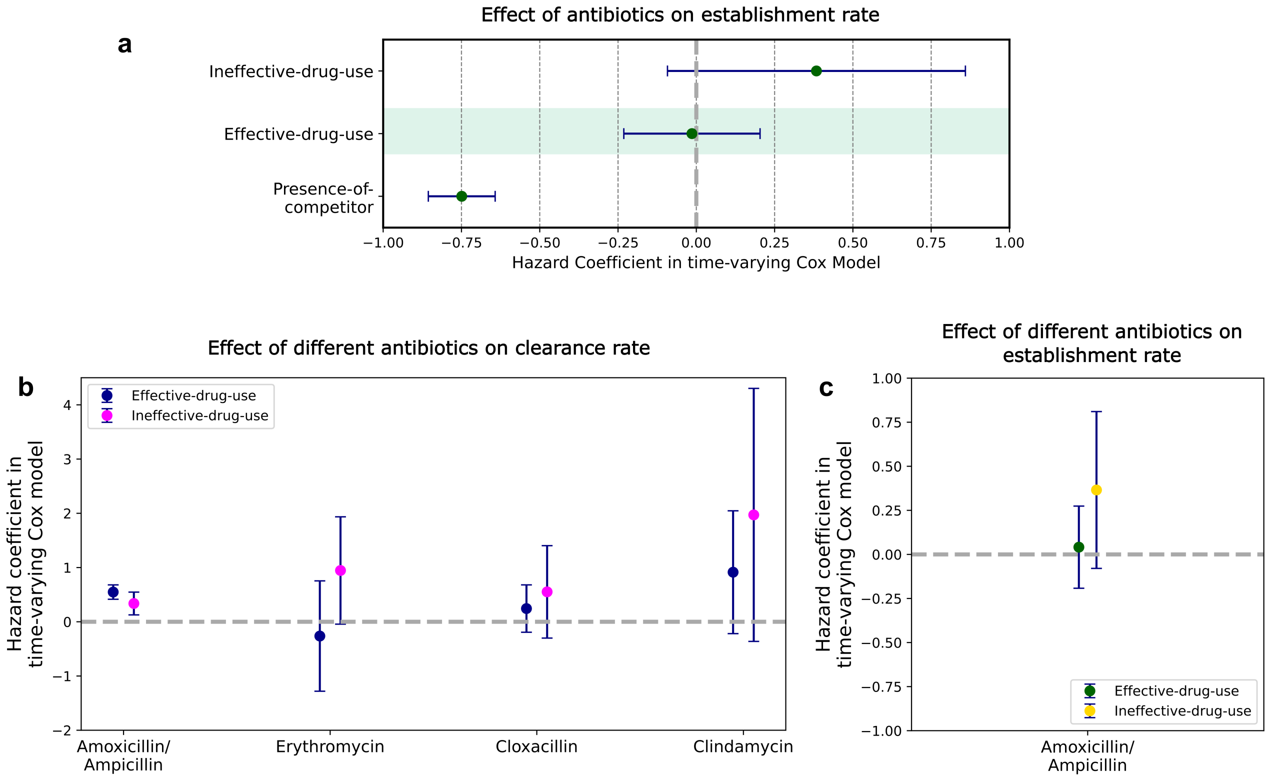


**Fig F.** Effects of drug consumption on clearance and establishment rates. (a) Effect of drug consumption on establishment rates across all antibiotics. The coefficient of *effective-drug-use* captures the effect of antibiotics on the establishment rate of drug-susceptible strains. (b,c) Effects of drug consumption on clearance rates (b), and establishment rates (c), analysed for each antibiotic in separate models. The hazard coefficients of *effective-drug-use* and *ineffective-drug-use* are shown for different antibiotics. For other drugs not shown in the figure, there is not enough data for convergence of the Cox model. The data underlying this figure can be found in https://doi.org/10.5281/zenodo.15800560

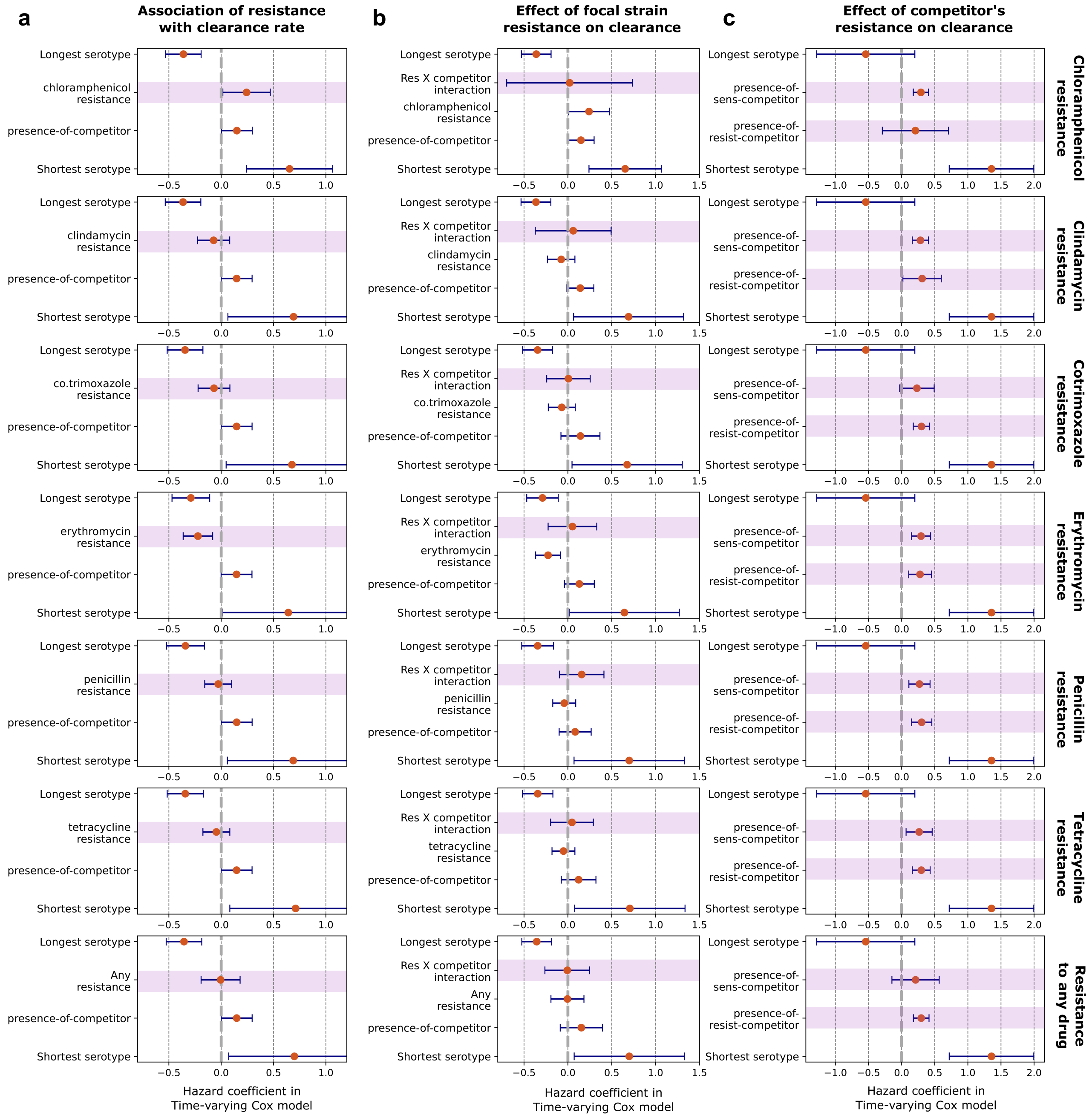


**Fig G.** Antibiotic-use-blind analyses of the effects of resistance on rate of clearance of serotypes (a) Does resistance to antibiotics affect the clearance rate of focal strain? The resistance variable tracks this for different antibiotics. (b) Does resistance to antibiotic affect clearance in the presence of a competitor? Here, the interaction term between resistance and presence-of-competitor tracks this effect for different drugs. (c) Does the resistance status of the co-coloniser strain affect clearance rate of the focal strain? We infer this based on the difference between coefficients of presence-of-sens-competitor and presence-of-resist-competitor for different drugs. The data underlying this figure can be found in https://doi.org/10.5281/zenodo.15800560

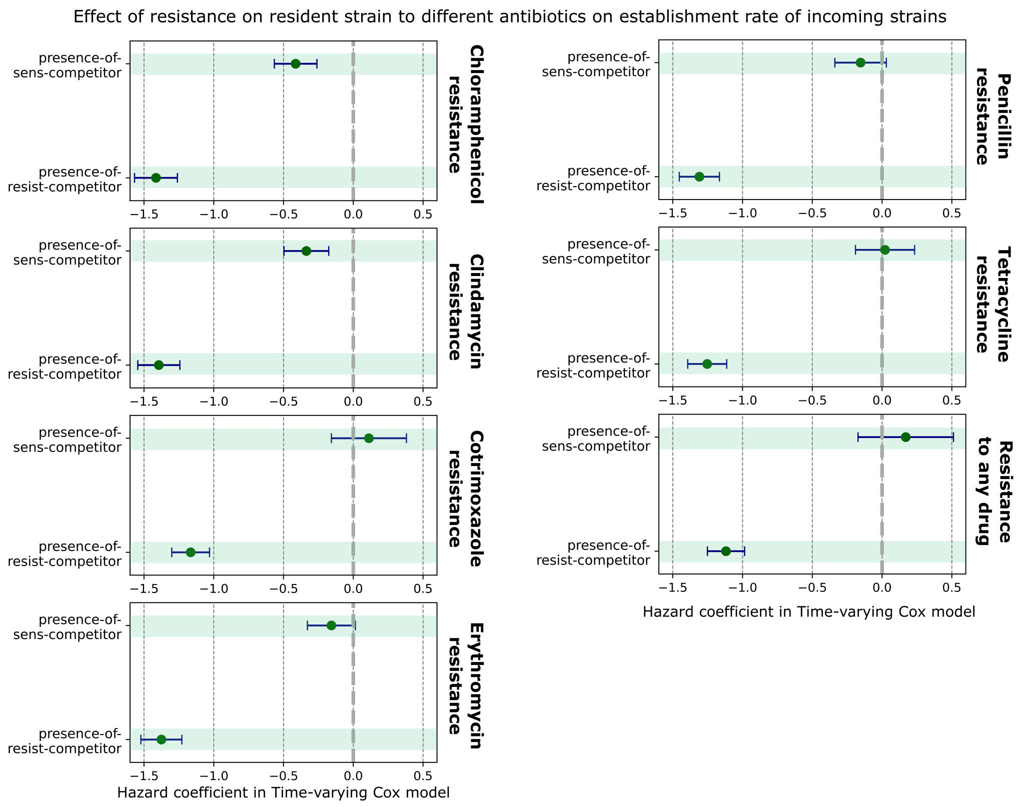


**Fig H.** Effects of resistance of within-host competitor on the establishment rate of focal strain, tested for resistance to different drugs. The difference in coefficients between presence-of-sens-competitor and presence-of-resist-competitor are shown for resistance to chloramphenicol, clindamycin, co-trimoxazole, erythromycin, penicillin, and tetracycline, and resistance to *any* drug. The data underlying this figure can be found in https://doi.org/10.5281/zenodo.15800560

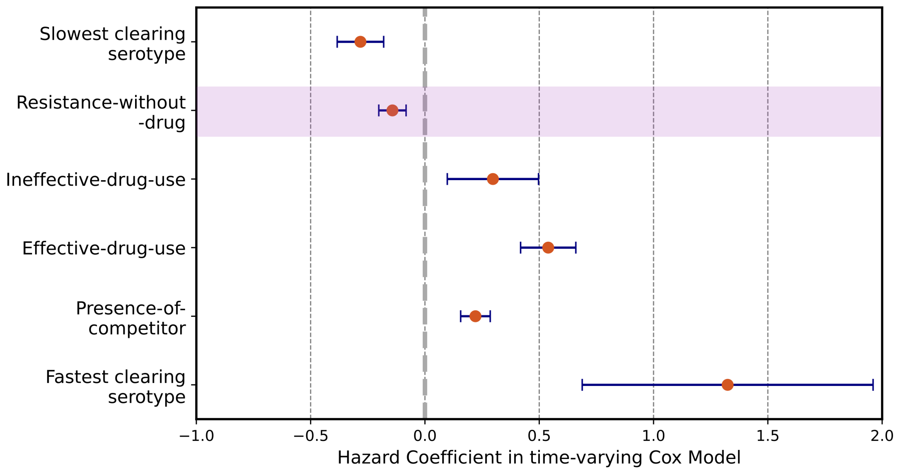


**Fig I.** Does antibiotic resistance affect clearance rate of strains? The covariate resistance-without-drug captures the effect of being resistant on rate of clearance. The negative coefficient indicates that resistant strains are associated with a lower clearance rate compared to susceptible strains. The data underlying this figure can be found in https://doi.org/10.5281/zenodo.15800560

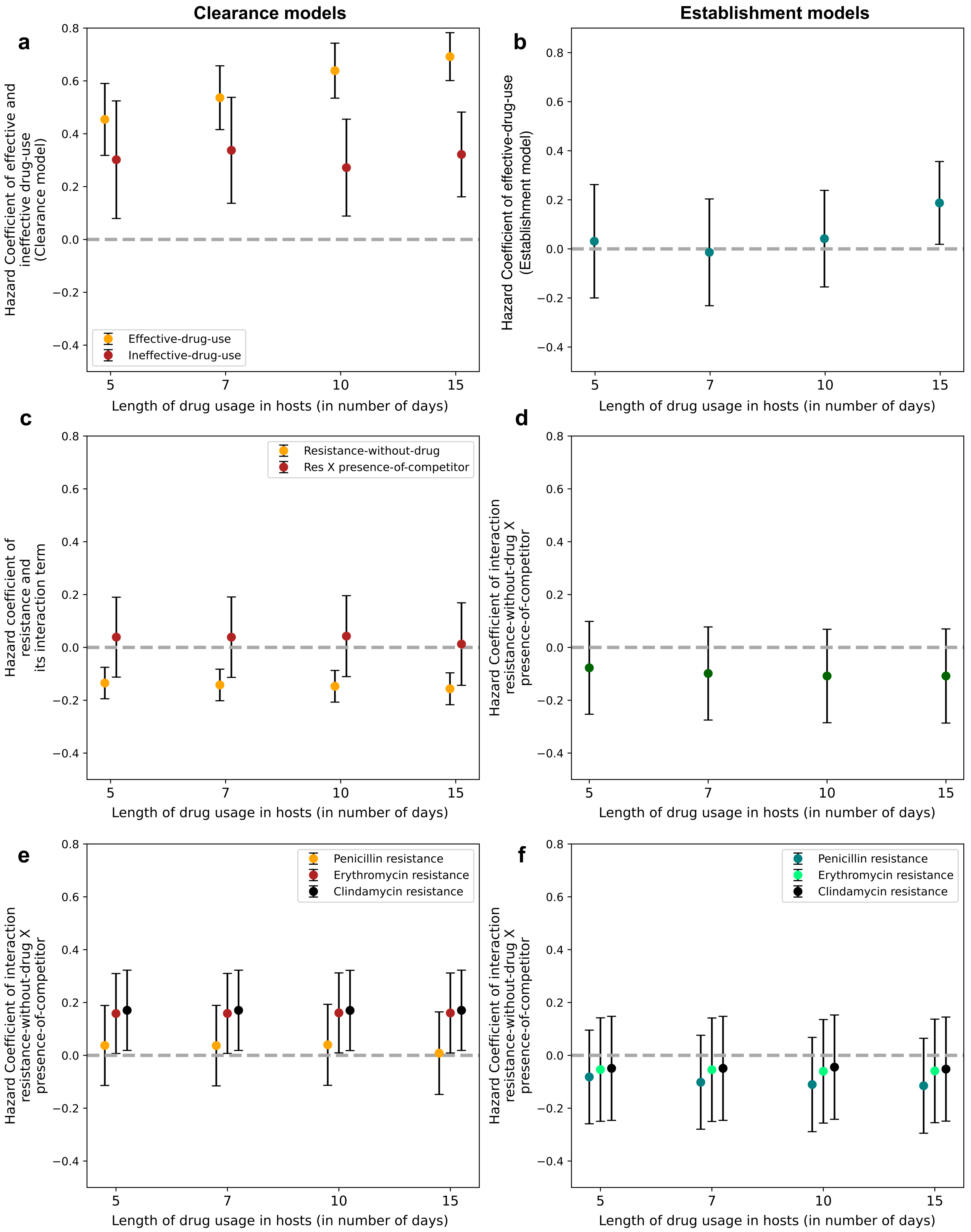


**Fig J.** Testing sensitivity of results to drug-use duration. a,c,e are based on models of time to clearance, and b,d,f are based on models of time to establishment. (a) The hazards coefficients of effective-drug-use and ineffective-drug-use on clearance rate are always positive. At 10 days and 15 days, the difference in hazards between effective-drug-use and ineffective-drug-use is significant. (b) The hazards coefficients of effective-drug-use and ineffective-drug-use on establishment rate are always non-significant, except at 15 days, when effective-drug-use has a positive coefficient. (c) The effect of resistance on clearance in the absence of drugs, and the effect of resistance on within-host competitive ability, are not sensitive to the durations of drug-use tested. (d) The within-cost of resistance during establishment is always negative and non-significant. (e,f) Within-host cost of resistance to different antibiotics during clearance (e) and establishment (f) are non-sensitive to duration of drug use. The data underlying this figure can be found in https://doi.org/10.5281/zenodo.15800560

Results on costs and benefits of antibiotic resistance using the subset of data where discordant resistance measurements are removed

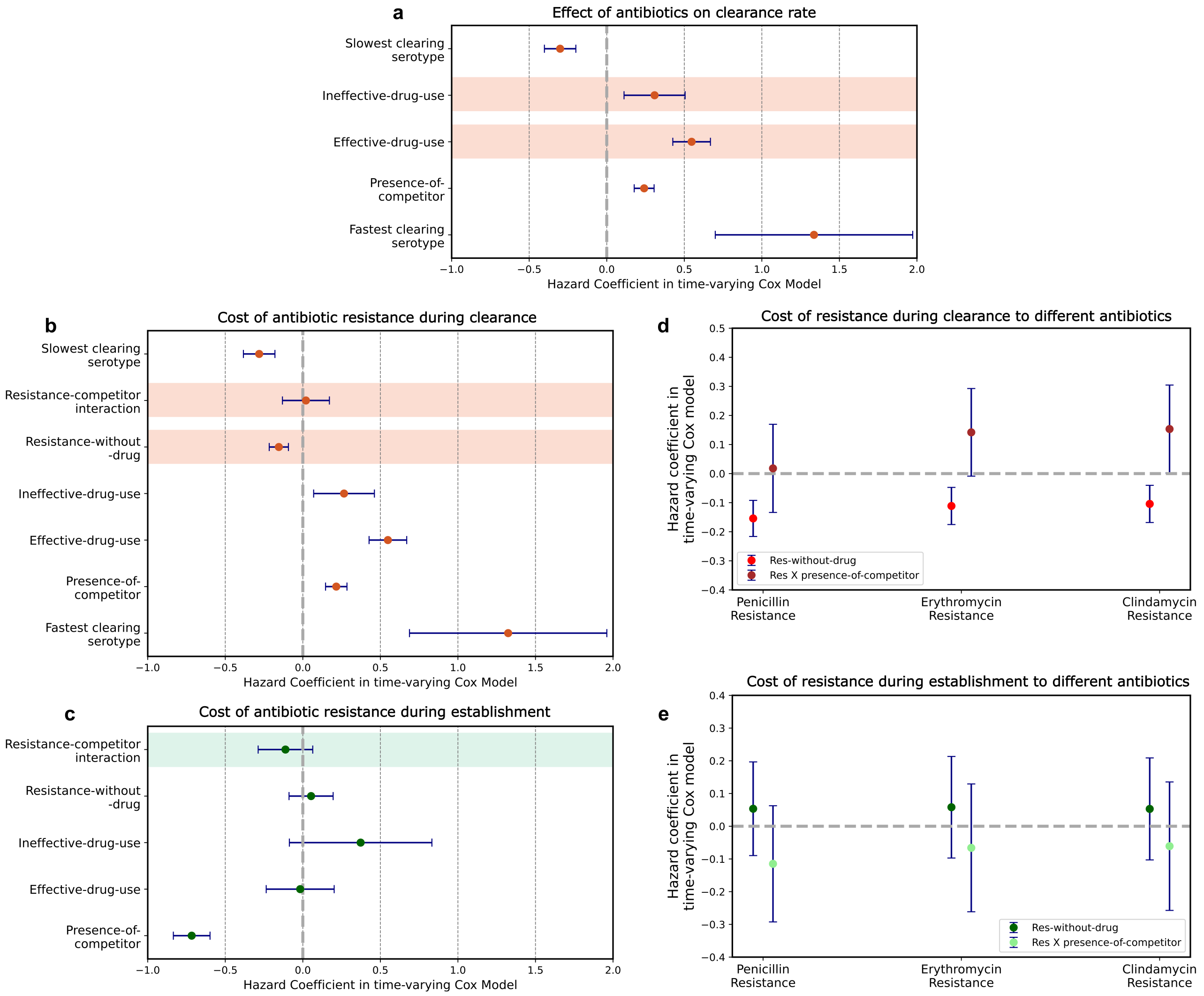


**Fig K.** Sensitivity of results to removing episodes of carriage where resistance status may have changed. a,b,d are models for clearance rate, while c,e show models for establishment rate. (a) The coefficients of *effective-drug-use* and *ineffective-drug-use*, capturing effect of antibiotic consumption on the clearance rate of resistant and susceptible strains, are positive, with a higher effect size on susceptible strains. (b) *Resistance-without-drug* has a negative hazard coefficient, indicating reduced clearance rates of resistant strains compared to susceptible strains. The interaction term here which captures the competitive cost of resistance during clearance has a positive non-significant hazard coefficient. (c) The interaction term that captures the competitive cost of being resistant during establishment into an occupied host niche is negative and non-significant. (d) The analysis in (b) is repeated for different resistances. The interaction terms corresponding to the cost of resistance during clearance to penicillin, erythromycin and clindamycin are positive. (e) The analysis in (c) is repeated for different resistances. The interaction terms corresponding to the cost of resistance during establishment to penicillin, erythromycin and clindamycin are negative. The data underlying this figure can be found in https://doi.org/10.5281/zenodo.15800560
